# A Variant-Resistant Linear Epitope in the SARS-CoV-2 Nucleocapsid Protein for Antigen Detection

**DOI:** 10.64898/2026.07.10.737673

**Authors:** Zhoujun Su, Junhao Guo, Hefeng Zhou, Jun Ni, Yongjun Cao, Liping Peng, Min Shao, Haitao Li

## Abstract

We previously generated a mouse monoclonal antibody, N179, against the SARS-CoV-2 nucleocapsid (N) protein and developed a colloidal gold-based immunochromatographic test strip. This assay achieved a detection limit of 2 ng/mL and displayed 98% concordance with RT-qPCR results. However, the precise epitope recognized by mAb N179 had not been defined. Using a panel of GST-fused N protein truncation fragments, we mapped the linear B-cell epitope recognized by mAb N179 to the flexible C-terminal tail of the N protein by Western blotting and ELISA. The minimal binding motif required for mAb N179 recognition was identified as ^390^QTVTLL^395^. Multiple sequence alignment of 11 representative SARS-CoV-2 lineages, including Alpha, Beta, Gamma, Delta, and Omicron subvariants BA.1, BA.2, and BA.3.2, revealed that this epitope was completely conserved across all variants analyzed. Stringent local pairwise alignment analysis using EMBOSS WATER further showed that the ^390^QTVTLL^395^ motif achieved a perfect 6/6 match exclusively in SARS-CoV-2; no identical sequence was detected in the seven common human coronaviruses, four influenza viruses, or five bat coronaviruses examined. Structural prediction analyses indicated that this region is surface-exposed and possesses a strong linear B-cell epitope propensity. Together, these findings identify ^390^QTVTLL^395^ as a specific molecular signature of SARS-CoV-2 among the viruses analyzed. Our results provide an epitope-level explanation for the sustained diagnostic reliability of the mAb N179-based assay against emerging variants, clarify the molecular basis for its lack of cross-reactivity, and may inform the rational design of SARS-CoV-2 diagnostics targeting conserved, mutation-resistant epitopes.

**Importance:** We identified the exact nucleocapsid protein epitope recognized by monoclonal antibody N179, a diagnostic antibody used in a colloidal gold rapid assay. The identified ^390^QTVTLL^395^ motif at residues 390 to 395 was conserved among the SARS-CoV-2 variants analyzed and was not present as an identical continuous sequence in the related respiratory viruses examined. This work supports precise epitope mapping as a useful strategy for evaluating and revalidating diagnostic antibodies as respiratory viruses evolve.

## Introduction

Rapid and accurate diagnostic testing remains essential for controlling transmission, enabling timely case identification, and guiding clinical management during the coronavirus disease 2019 (COVID-19) pandemic caused by severe acute respiratory syndrome coronavirus 2 (SARS-CoV-2) (1). Reverse transcription-quantitative PCR (RT-qPCR) is still regarded as the reference standard. However, the requirement for sophisticated laboratory infrastructure and trained personnel, coupled with prolonged turnaround times, restricts its application in point-of-care testing and large-scale screening settings (2, 3). Antigen rapid diagnostic tests (Ag-RDTs) based on lateral flow immunochromatography therefore provide a practical alternative owing to their operational simplicity, rapid turnaround time, and low cost (4, 5).

The nucleocapsid (N) protein is one of the four major structural proteins of SARS-CoV-2 (6). Because it is abundantly expressed in infected cells, highly immunogenic, and relatively conserved across viral lineages, the N protein serves as a major target for antigen detection (7–9). Nevertheless, the continued emergence of SARS-CoV-2 variants threatens the reliability of antibody-based diagnostic assays (10–13). Mutations in the N protein, particularly those located within antibody-recognition epitopes, can diminish or even abolish antibody binding, thereby resulting in false-negative outcomes (12). For instance, the D399N substitution in the B.1.429 variant substantially compromised the performance of the Quidel Sofia SARS Antigen FIA test, demonstrating that a single amino acid substitution can impair diagnostic sensitivity (14). Similarly, antibodies targeting the N-terminal RNA-binding domain or the central dimerization domain can be disrupted by mutations within their respective epitopes (15). These observations underscore the critical need to identify and characterize monoclonal antibodies that recognize highly conserved linear epitopes in the N protein (14, 15).

We previously generated a mouse monoclonal antibody, N179, against the SARS-CoV-2 N protein and used it to develop a colloidal gold-based immunochromatographic test strip with a detection limit of 2 ng/mL and 98% concordance with RT-qPCR results (16). Despite the diagnostic performance of this strip, the precise epitope recognized by mAb N179 remained undefined. Several critical questions thus remained unresolved: whether the epitope is conserved across variants, whether it is unique among related respiratory viruses and what molecular features underlie mAb N179 recognition. In this study, we fine-mapped the linear B-cell epitope recognized by mAb N179 using Western blotting and ELISA with a panel of GST-fused N protein truncation fragments. We further assessed the conservation of this epitope across representative SARS-CoV-2 variants, examined its sequence specificity among seven human coronaviruses, four influenza viruses, and five bat coronaviruses, and characterized its predicted structural features. Together, these analyses clarify the epitope-level basis of mAb N179 recognition, evaluate the potential of this epitope motif as a robust diagnostic target, and provide a molecular explanation for the sustained specificity and lack of cross-reactivity observed with the mAb N179-based assay.

## Materials and Methods

### Expression and purification of full-length N protein and antibody validation

The recombinant plasmid pET-28a(+)-CorN was transformed into *Escherichia coli* BL21(DE3) competent cells (Cat.No.BC101; Biomed). Protein expression was induced with 0.5 mM isopropyl-β-D-thiogalactopyranoside (IPTG) at 16°C for 16 h. The cells were harvested, resuspended in lysis buffer containing 50 mM Tris-HCl, 500 mM NaCl, 10 mM imidazole, and 1 mM phenylmethylsulfonyl fluoride (PMSF) at pH 7.4, and disrupted by ultrasonication at 200 W, using 3-s pulses with 5-s intervals for a total of 20 min. The lysate was centrifuged at 12,000 × g for 30 min at 4°C, and the supernatant was collected for subsequent purification. The His-tagged recombinant N protein (CorN) in the supernatant was purified using an ÄKTA purification system equipped with a HisTrap Excel column (Cat. No. 17-3712-06; GE Healthcare). The column was pre-equilibrated with binding buffer, which had the same composition as the lysis buffer. After sample loading, the column was washed with binding buffer to remove unbound and nonspecifically bound proteins. The target protein was eluted over 10 column volumes with a linear imidazole gradient from 10 to 500 mM generated by mixing binding buffer (50 mM Tris-HCl, 500 mM NaCl, 10 mM imidazole, pH 7.4)with elution buffer (50 mM Tris-HCl, 500 mM NaCl, 500 mM imidazole, pH 7.4). Protein elution was monitored by absorbance at 280 nm (A₂₈₀), and peak fractions were collected. The eluted protein was concentrated and buffer-exchanged into phosphate-buffered saline (PBS; pH 7.4) using an ultrafiltration device with a 10 kDa molecular weight cutoff (Cat. No. UFC9010; Millipore). The purified protein was quantified using a bicinchoninic acid (BCA) assay and analyzed by sodium dodecyl sulfate-polyacrylamide gel electrophoresis (SDS-PAGE), followed by Western blotting to verify its specific reactivity with mAb N179. Briefly, protein samples were denatured at 100°C for 10 min, separated on a 12% SDS-PAGE gel, and transferred onto a polyvinylidene difluoride (PVDF) membrane (Cat. No. ISEQ00010; Millipore). The membrane was blocked with 5% nonfat milk in Tris-buffered saline with Tween 20 (TBST) for 2 h at room temperature and then incubated overnight at 4°C with mAb N179 as the primary antibody (IgG2a/κ; 1:10,000 dilution in TBST; sequencing data are provided in supplementary material Table S1). After washing with TBST, the membrane was incubated with horseradish peroxidase (HRP)-conjugated goat anti-mouse IgG secondary antibody diluted 1:2,000 in TBST for 1 h at room temperature. Following additional washes, chemiluminescent signals were detected using an enhanced chemiluminescence (ECL) kit (Cat. No. P0018AS; Beyotime).

### Construction and expression of truncated GST fusion proteins

To localize the region recognized by mAb N179, primers were designed based on the SARS-CoV-2 reference genome sequence in GenBank (accession no. NC_045512.2; Table 1), and a series of recombinant plasmids expressing GST-fused full-length N protein or N protein truncated fragments covering different C-terminal regions was constructed. Briefly, the pGEX-4T-3 vector was linearized by double digestion with BamHI and NotI (Cat. No. 10133983 and 10133990; NEB). Each truncated fragment was amplified by polymerase chain reaction (PCR). The fragments marked with an asterisk in Table 1 were cloned into the linearized pGEX-4T-3 vector using T4 DNA ligase, whereas the remaining fragments were cloned using a homologous recombination kit (Cat No. C117; Vazyme). All constructs were verified by Sanger sequencing.

**Table 1.**
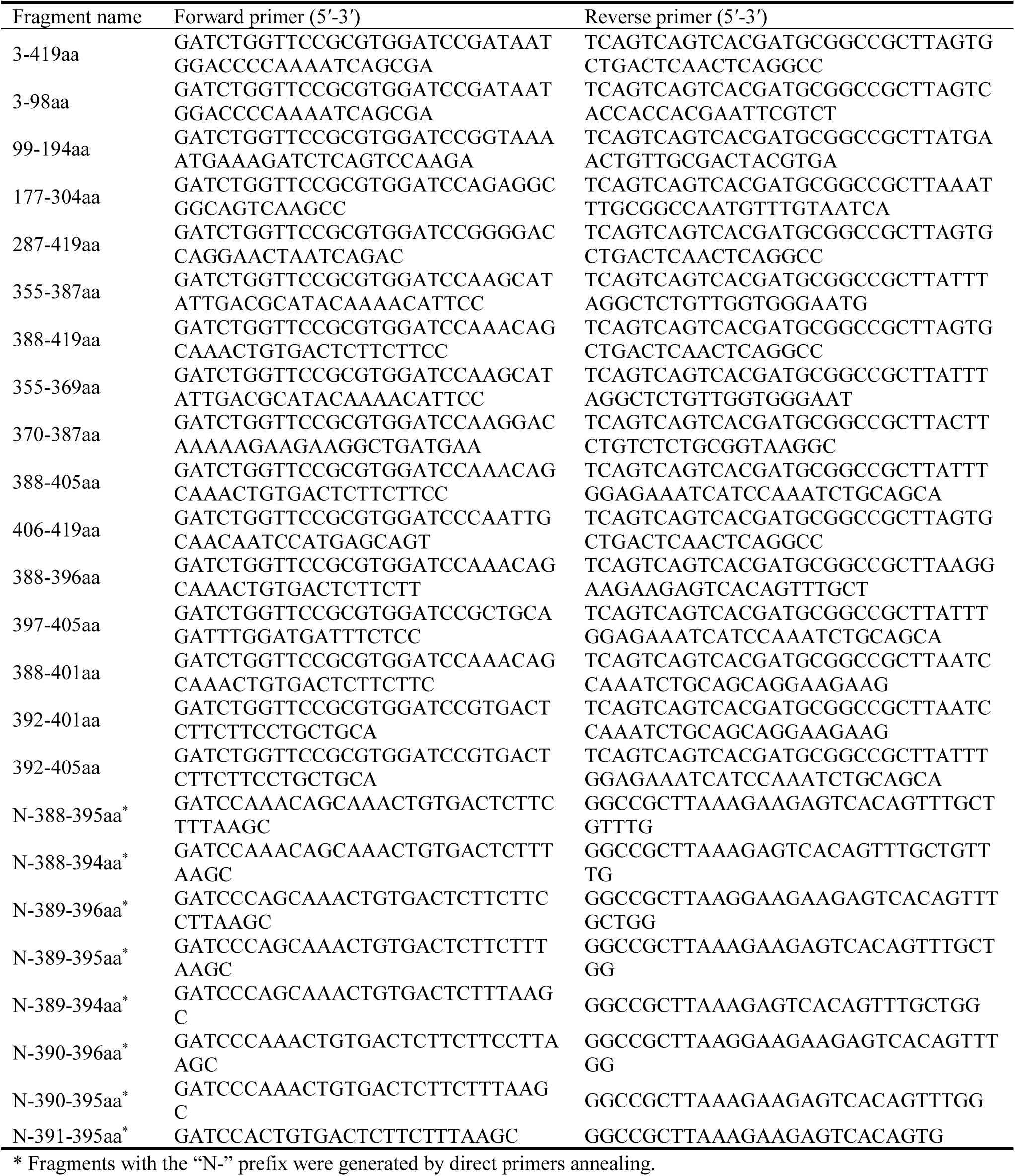
PCR primers used to amplify the SARS-CoV-2 N protein gene and truncated fragments.

For protein expression, each recombinant plasmid was transformed into Escherichia coli BL21(DE3) competent cells (Cat. No. BC101; Biomed). Single colonies were picked, and inoculated into Luria-Bertani (LB) medium containing ampicillin and cultured overnight at 37°C with shaking at 250 rpm. The overnight cultures were diluted 1:100 into 200 mL of fresh LB medium containing ampicillin and grown at 37°C with shaking at 250 rpm until the optical density at 600 nm (OD₆₀₀) reached 0.5. IPTG was then added to a final concentration of 0.5 mM, and protein expression was induced at 16°C for 16 h with shaking at 180 rpm. The cells were harvested by centrifugation at 10,000 × g for 10 min at 4°C.

The cell pellets were resuspended in PBS (pH 7.4) containing 1 mM PMSF and disrupted by ultrasonication at 200 W, using 3-s pulse with 5-s intervals for a total of 30 min. The lysates were centrifuged at 12,000 × g for 30 min at 4°C, and the supernatants were collected for subsequent purification. Glutathione S-transferase (GST)-tagged truncated proteins in the supernatant were purified using an ÄKTA purification system equipped with a GST affinity column (Cat. No. SA010C55; Smart-Lifesciences). The column was pre-equilibrated with PBS (pH 7.4). After sample loading, the column was washed with PBS (pH 7.4) to remove unbound proteins, and the target proteins were eluted with elution buffer containing 50 mM Tris-HCl and 10 mM reduced glutathione at pH 8.0. Protein elution was monitored by absorbance at 280 nm (A₂₈₀), and peak fractions were collected. The eluted proteins were concentrated and buffer-exchanged into PBS (pH 7.4) using an ultrafiltration device with a 3 kDa molecular weight cutoff (Cat. No. UFC9003; Millipore) to remove reduced glutathione.

### Epitope screening by Western blotting and ELISA

Purified GST-tagged truncated proteins were separated by 12% SDS-PAGE and transferred onto PVDF membranes. Blocking, antibody incubation, washing, and chemiluminescence detection were performed as described above for Western blot analysis of full-length N protein.

For enzyme-linked immunosorbent assay (ELISA), purified GST-tagged truncated proteins were quantified using a bicinchoninic acid (BCA) protein assay kit (Cat. No. P0012, Beyotime), diluted to 1 μg/mL in coating buffer consisting of 0.05 M carbonate-bicarbonate buffer (pH 9.6), and added to 96-well plates at 100 μL per well. The plates were coated overnight at 4°C. After blocking with TBST containing 5% bovine serum albumin (BSA) for 2 h at 4°C and washing three times with TBST, mAb N179 as the primary antibody at 1:10,000 dilution was added and incubated at 37°C for 1 h. Following three washes with TBST, HRP-conjugated goat anti-mouse IgG secondary antibody at 1:2,000 dilution was added and incubated at 37°C for 45 min. After three additional washes with TBST, 100 μL of 3,3’-tetramethylbenzidine (TMB) substrate solution (Cat. No. P0210; Beyotime) was added to each well, and the plates were incubated at 37°C for 15 min in the dark. The reaction was stopped by adding ELISA stop solution (Cat. No. P0215; Beyotime), and the optical density at 450 nm (OD₄₅₀) was measured. A sample was considered positive when the ratio of its OD₄₅₀ value to that of the negative control (P/N) was ≥2.1. GST protein expressed from the empty vector was used as the negative control.

### Sequence retrieval and multiple sequence alignment

To assess epitope conservation, N protein sequences from 11 representative SARS-CoV-2 lineages, including Alpha, Beta, Gamma, Delta, and Omicron BA.1, BA.2, and BA.3.2, were retrieved from the National Center for Biotechnology Information (NCBI) GenBank database (accessed 15 February 2026). To evaluate the sequence specificity of the ^390^QTVTLL^395^ epitope among related viruses, nucleoprotein sequences were obtained from the NCBI Protein database for the following viruses: seven human coronaviruses, including SARS-CoV-2 (NC_045512.2), SARS-CoV (NC_004718.3), Middle East respiratory syndrome coronavirus (MERS-CoV; NC_019843.3), human coronavirus OC43 (HCoV-OC43; NC_005147.1), HCoV-229E (NC_002645.1), HCoV-NL63 (NC_005831.1), and HCoV-HKU1 (NC_006577.2); four influenza virus nucleoproteins (NPs), including influenza A virus H1N1 (NP_040982), influenza A virus H3N2 (P22435.2), influenza B virus Victoria lineage (AGX24738), and influenza B virus Yamagata lineage (AAA82969); and five bat coronaviruses, including GCCDC1 (NC_030886), HKU9-1 (NC_009021), CMR704-P12 (NC_048212), BM48-31 (NC_014470), and CDPHE15 (NC_022103). Multiple sequence alignments were performed using Clustal Omega (https://www.genome.jp/tools-bin/clustalw) with default parameters, and the results were visualized with ESPript 3.2 (https://espript.ibcp.fr/ESPript/ESPript/index.php).

For stringent local pairwise alignment, the EMBOSS WATER algorithm was used with the position-specific amino acid matrix (PAM) 30 substitution matrix (https://www.ebi.ac.uk/jdispatcher/psa/emboss_water), a gap opening penalty of 10, and a gap extension penalty of 0.5 to evaluate the sequence matches of the ^390^QTVTLL^395^ motif across viral nucleoproteins.

### Structural predictions

The secondary structure of the ^390^QTVTLL^395^ epitope was predicted using PSIPRED (https://bioinf.cs.ucl.ac.uk/psipred), and its hydrophobicity was evaluated with ProtScale using the Kyte-Doolittle hydropathy scale(https://web.expasy.org/protscale/) with a window size of 9. Linear B-cell epitope propensity was assessed using BepiPred 3.0 (https://services.healthtech.dtu.dk/services/BepiPred-3.0/). Surface exposure of the ^390^QTVTLL^395^ region was analyzed using the AlphaFold 3-predicted structural model of the full-length N protein (AlphaFold Database accession no. AF-0000000365840321-v1, corresponding to Universal Protein Resource (UniProt) P0DTC9; accessed 8 April 2026; https://alphafold.ebi.ac.uk/entry/AF-0000000365840321). Panels A to D in Fig 7 were generated from the outputs of PSIPRED, ProtScale, and ScanProsite (https://prosite.expasy.org/scanprosite/) using a custom Python script. Panel E was a direct screenshot of the BepiPred 3.0 web server output, Panel F was visualized with PyMOL version 3.2.0a.

### Statistical analysis

ELISA experiments were performed independently three times, with three replicate wells included in each experiment. Data are presented as the mean ± standard deviation (SD). Samples were considered positive when the P/N ratio was ≥2.1 as described above. GraphPad Prism 8 (GraphPad Software, San Diego, CA, USA) was used for statistical analysis and graph preparation.

## Results

### Validation of mAb N179 Recognition of Recombinant N Protein

To verify the binding specificity of mAb N179 for recombinant N protein, we expressed and purified His-tagged recombinant N protein (CorN) and analyzed it by Western blotting. SDS-PAGE analysis showed a distinct protein band at approximately 50 kDa following IPTG induction, corresponding to the expected molecular weight of CorN. This band was detected mainly in the supernatant fraction, indicating that CorN was expressed as a predominantly soluble protein (Fig 2A). Western blotting further confirmed that mAb N179 specifically recognized the purified CorN, yielding a single immunoreactive band at the expected molecular weight (Fig 2B).

**Fig 1.**
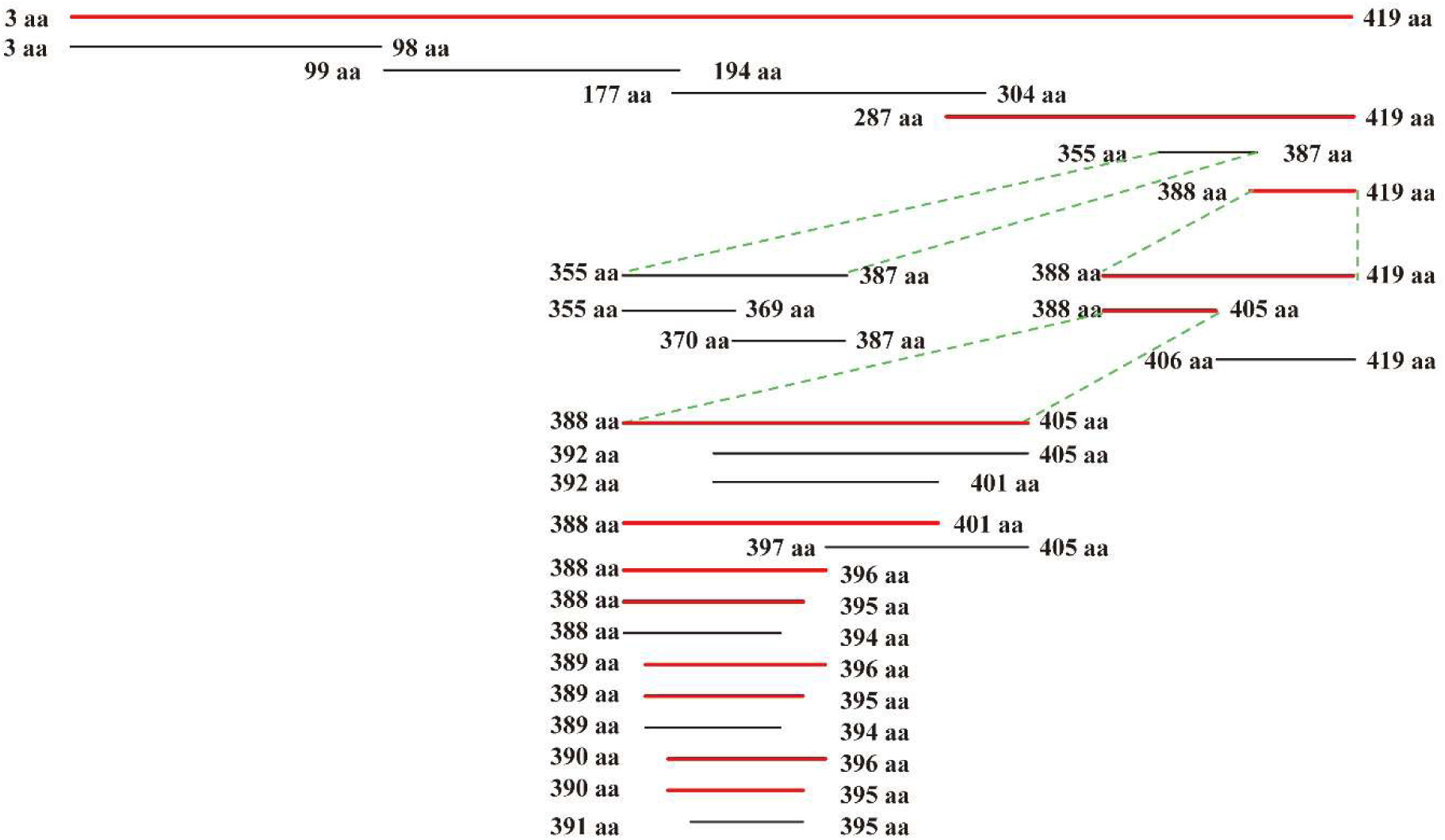
Schematic representation of the truncated SARS-CoV-2 N protein fragments used for epitope mapping. Amino acid positions are indicated for each fragment. The near-full-length N protein fragment (3–419 aa) and progressively truncated fragments were analyzed by Western blotting and ELISA. Red bars indicate fragments that showed positive reactivity with mAb N179.

**Fig 2.**
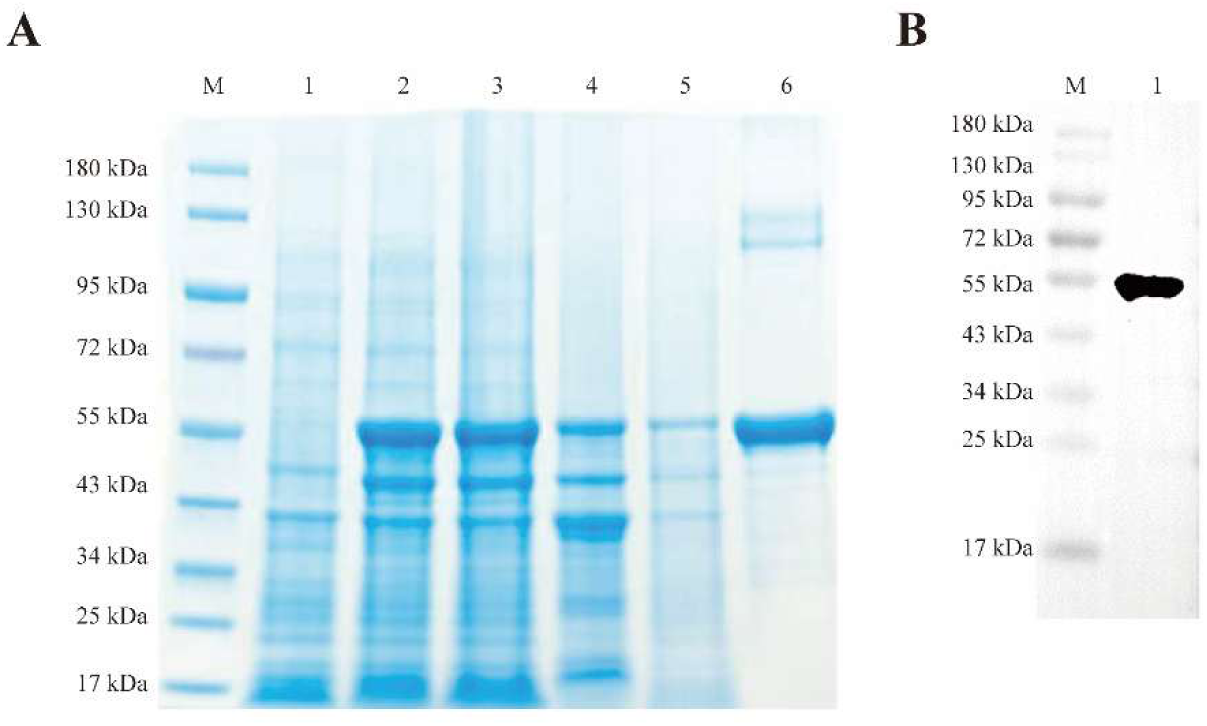
Expression, purification, and Western blot validation of recombinant SARS-CoV-2 N protein. (A) SDS-PAGE analysis of recombinant N protein expression and purification. Lane M, protein molecular weight marker; lane 1, whole-cell lysate before induction; lane 2, whole-cell lysate after IPTG induction; lane 3, supernatant fraction after ultrasonication; lane 4, pellet fraction after ultrasonication; lane 5, flow-through from the HisTrap Excel affinity column; lane 6, purified recombinant N protein after elution. (B) Western blot analysis of purified recombinant N protein probed with mAb N179. Lane M, protein molecular weight marker; lane 1, purified recombinant N protein.

### Minimal Binding Sequence of mAb N179 recognition is Residue 390-395

To localize the epitope recognized by mAb N179, we expressed a series of GST-fused truncated proteins spanning the full-length N protein and selected regions of its C-terminal portion. SDS-PAGE analysis showed that each truncated protein migrated as a distinct band at the expected molecular weight following IPTG induction, indicating successful expression of all constructs (Fig 3A).

**Fig 3.**
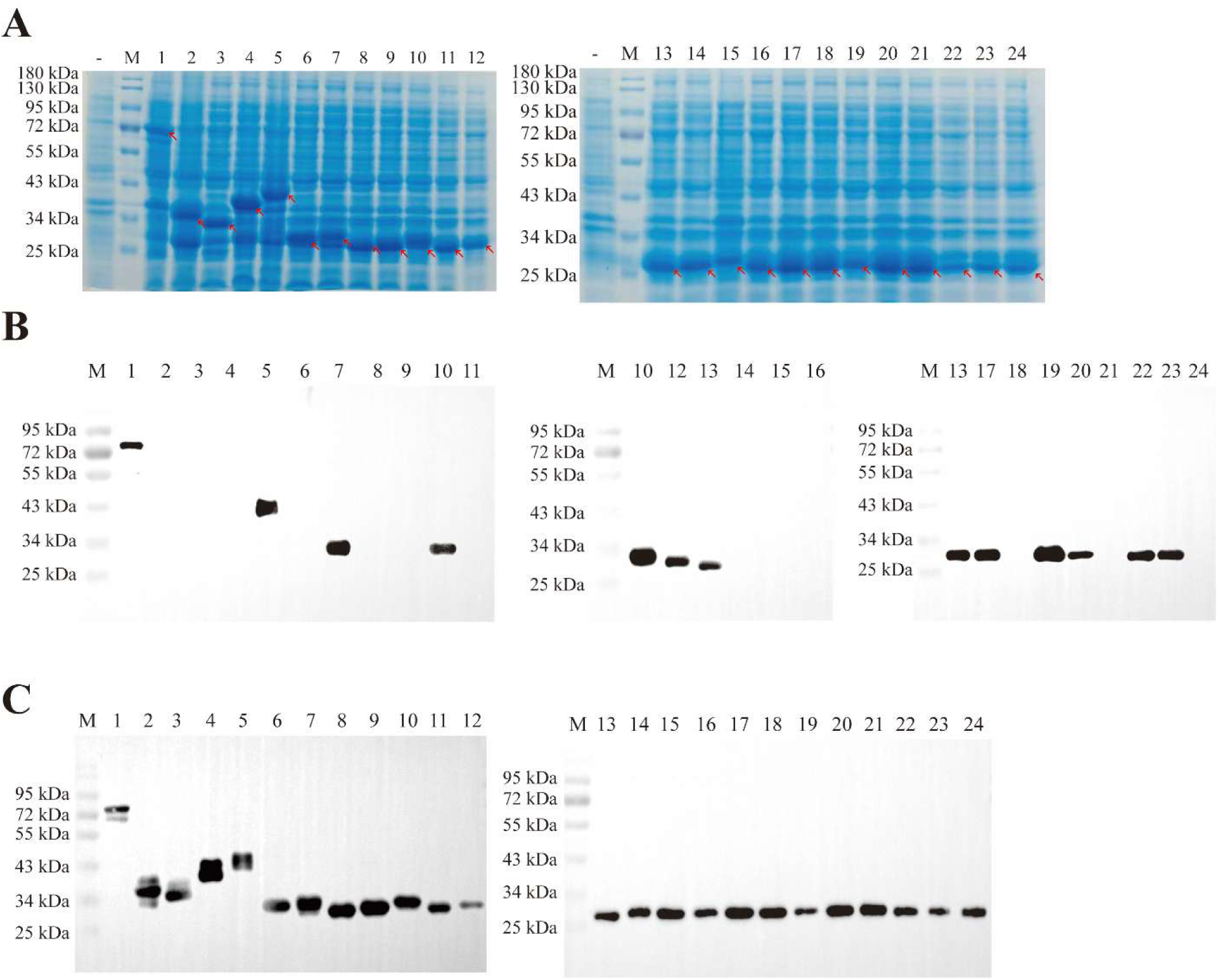
Western blot mapping of the antigenic epitope recognized by mAb N179. (A) Coomassie blue-stained SDS-PAGE gel showing the expression of each GST-tagged truncated protein fragments. Lane “−”, whole-cell lysate before induction; (B) Western blot analysis of the reactivity of the truncated fragments with mAb N179. Lane M, protein molecular weight marker; lane 1, 3-419 aa; lane 2, 3-98 aa; lane 3, 99-194 aa; lane 4, 177-304 aa; lane 5, 287-419 aa; lane 6, 355-387 aa; lane 7, 388 −419 aa; lane 8, 355- 369 aa; lane 9, 370-387 aa; lane 10, 388-405 aa; lane 11, 406-419 aa; lane 12, 388-401 aa; lane 13, 388-396 aa; lane 14, 397-405 aa; lane 15, 392-405 aa; lane 16, 392-401 aa; lane 17, 388-395 aa; lane 18, 388-394 aa; lane 19, 389-396 aa; lane 20, 389-395 aa; lane 21, 389-394 aa; lane 22, 390-396 aa; lane 23, 390-395 aa; lane 24, 391- 395 aa. The smallest fragment producing a positive signal was 390-395 aa. (C) Western blot analysis using an anti-GST antibody as a loading control, confirming the expression and loading of each truncated protein.

Western blotting showed that mAb N179 recognized the full-length N protein (residues 3–419) and the C-terminal fragments 287–419 aa, 355–387 aa, and 388–419 aa, but did not react with the N-terminal fragment (residues 3–98) or the central-region fragments 99–194 aa and 177–304 aa. These results initially localized the epitope to the C-terminal region (residues 388–419) (Fig 3B, lanes 1–11). Further truncation narrowed the reactive region to residues 388–396 (Fig 3B, lanes 12–16). Sequential single-residue truncation demonstrated that deletion of either glutamine at position 390 (Q390; lane 21) or leucine at position 395 (L395; lane 24) abolished immunoreactivity. Thus, the minimal binding motif required for mAb N179 recognition was defined as ^390^QTVTLL^395^ (lane 23). Western blotting with an anti-GST monoclonal antibody further confirmed that all truncated proteins carried the GST tag (Fig 3C, lanes 1–24).

To quantitatively validate the epitope-mapping results, the same set of GST-tagged truncated proteins was purified by GST affinity chromatography (Fig 4A), quantified by BCA assay, and coated onto 96-well plates at 1 μg/mL for ELISA analysis (Fig 4B). Consistent with the Western blotting data, mAb N179 reacted strongly with fragments spanning the 388–419 aa region, including 388–419 aa, 388–405 aa, 388–396 aa, 392–405 aa, and 392–401 aa, with all OD₄₅₀ values exceeding the positivity threshold (P/N ≥ 2.1). In contrast, fragments lacking the ^390^QTVTLL^395^ core motif, such as 397–405 aa and 406–419 aa, displayed OD₄₅₀ values below the threshold. In the fine-mapping analysis, only the fragment retaining the intact ^390^QTVTLL^395^ motif (residues 390–395) produced a strong positive signal, whereas fragments missing a single residue at either end, such as 389–394 aa or 391–395 aa, yielded markedly reduced OD₄₅₀ values comparable to that of the GST control. These quantitative ELISA results were consistent with the Western blotting findings, further confirmed that ^390^QTVTLL^395^ is required for recognition by mAb N179 (Fig 4C).

**Fig 4.**
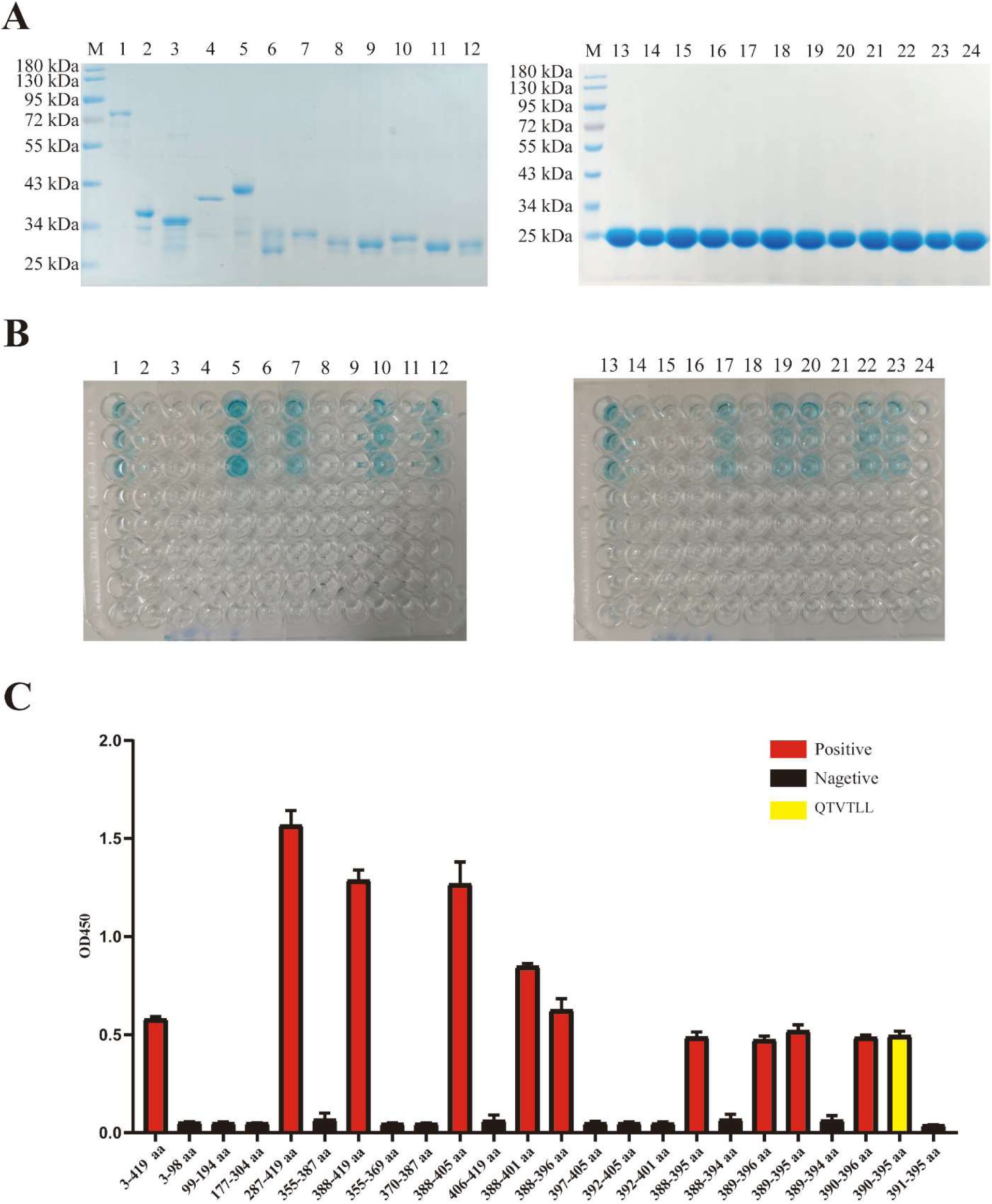
ELISA validation of the antigenic epitope recognized by mAb N179. (A) SDS-PAGE analysis of the purified GST-fused truncated proteins fragments. (B) ELISA color development with TMB substrate for each truncated fragment. (C) OD₄₅₀ values for each truncated fragment after reaction with mAb N179. Fragment numbering is consistent with that in Fig. 3B. OD₄₅₀ values are presented as the mean ± SD from three independent experiments; the dashed line indicates the positivity threshold (P/N ≥ 2.1).

### The QTVTLL Sequence is Complete Conserved in Major SARS-CoV-2 Lineages Analyzed

To evaluate the conservation of the ^390^QTVTLL^395^ epitope among circulating variants, we performed multiple sequence alignment of N protein sequences from 11 representative SARS-CoV-2 lineages. The results revealed that this epitope was completely conserved across all variants analyzed, including Alpha, Beta, Gamma, and Delta, as well as Omicron BA.1, BA.2, and BA.3.2 sub-lineages, with no amino acid substitutions detected in this region (Fig 5).

**Fig 5.**
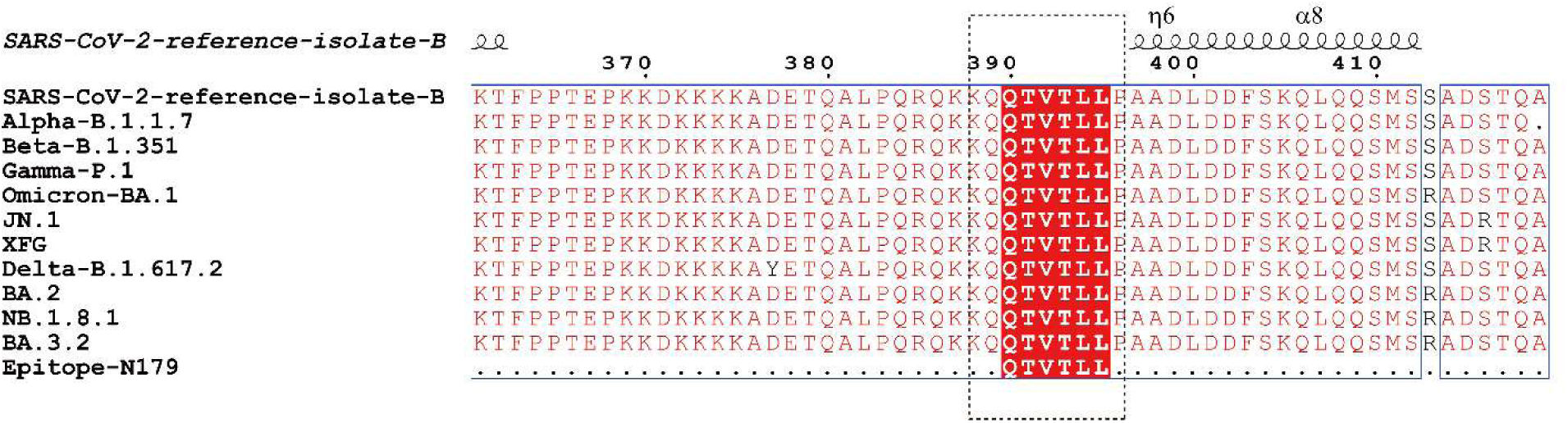
Conservation analysis of the 390QTVTLL395 epitope among representative SARS-CoV-2 variants. Multiple sequence alignment of N protein sequences from 11 representative SARS-CoV-2 lineages was performed using Clustal Omega and visualized with ESPript 3.2. The ^390^QTVTLL^395^ region is highlighted with a dashed box. The lineages analyzed include Alpha, Beta, Gamma, Delta, and Omicron sub-lineages BA.1, BA.2 and BA.3.2. Secondary structure predictions are shown above the sequences with helix and coil symbols, indicating this region adopts a predicted coil/random-coil conformation.

### The QTVTLL Sequence in SARS-CoV-2 is Exclusive Among Analyzed Variants

To evaluate the sequence specificity of the ^390^QTVTLL^395^ epitope among related viruses, we aligned the N protein sequences of seven human coronaviruses, the NP sequences of four influenza viruses, and the N proteins sequences of five bat coronaviruses. No exact match to the ^390^QTVTLL^395^ epitope was identified in any of the 16 non-SARS-CoV-2 viruses analyzed (Fig 6). SARS-CoV harbored a single amino acid substitution, Q390P (glutamine at position 390 to proline), in this region, whereas the bat coronavirus BM48-31 contained a Q390A (glutamine at position 390 to alanine) substitution. The corresponding regions in MERS-CoV and the four seasonal human coronaviruses, HCoV-OC43, HCoV-229E, HCoV-NL63, and HCoV-HKU1, differed substantially from the SARS-CoV-2 sequence and retained only two to three matching residues. Similarly, only two to three scattered matching residues were detected in the NP proteins sequences of the four influenza viruses and the N proteins sequences of the remaining bat coronaviruses, and no continuous homologous motif was observed (Fig 6).

**Fig 6.**
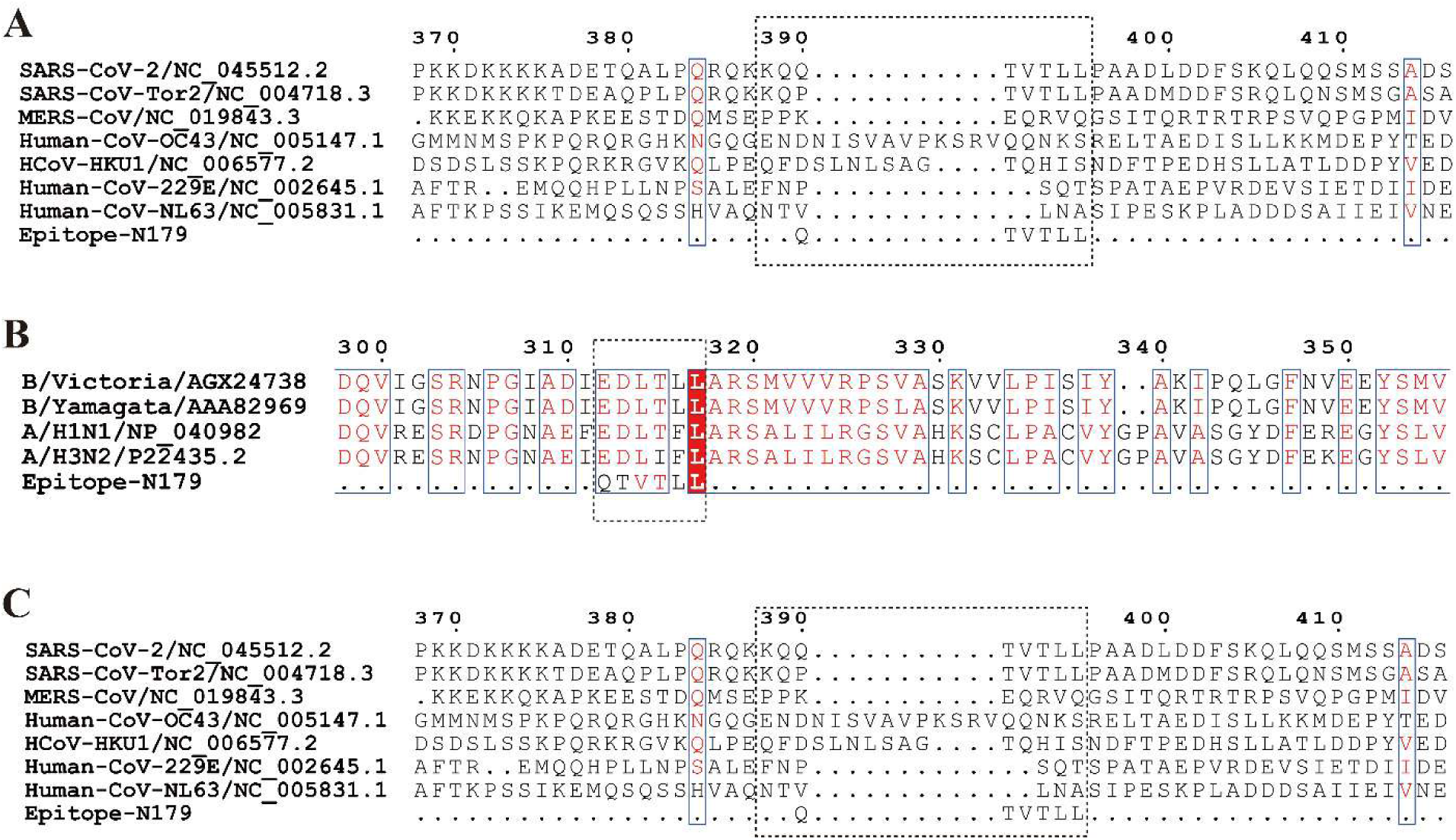
Cross-species conservation analysis of the 390QTVTLL395 epitope among human coronaviruses, influenza viruses, and bat coronaviruses. Multiple sequence alignment of the N or NP protein regions from seven human coronaviruses, four influenza viruses, and five bat coronaviruses was performed using Clustal Omega and visualized with ESPript 3.2. The ^390^QTVTLL^395^ epitope is highlighted with a dashed box. The motif was fully matched exclusively in SARS-CoV-2; the most closely related viruses in this analysis, SARS-CoV and BM48-31, each contained a single residue substitution resulting in a 5/6 match, whereas no continuous matched motif was present in any of the other species analyzed.

Stringent local pairwise alignment using EMBOSS WATER algorithm demonstrated that the ^390^QTVTLL^395^ motif achieved an exact 6/6 match only in SARS-CoV-2, while the closest related coronaviruses retained a single mismatched residue (Table 2). SARS-CoV and the bat coronavirus BM48-31 each carried a single amino acid substitution, Q390P and Q390A respectively, resulting in a 5/6 match (83.3% identity). The remaining 13 viruses matched only two to three residues in their best-matching regions corresponding to 33.3% to 50.0% identity, and none contained a continuous six-residue motif matching ^390^QTVTLL^395^ (Table 2). These results support the identification of ^390^QTVTLL^395^ as a SARS-CoV-2-specific linear motif among the viruses analyzed.

**Table 2.**
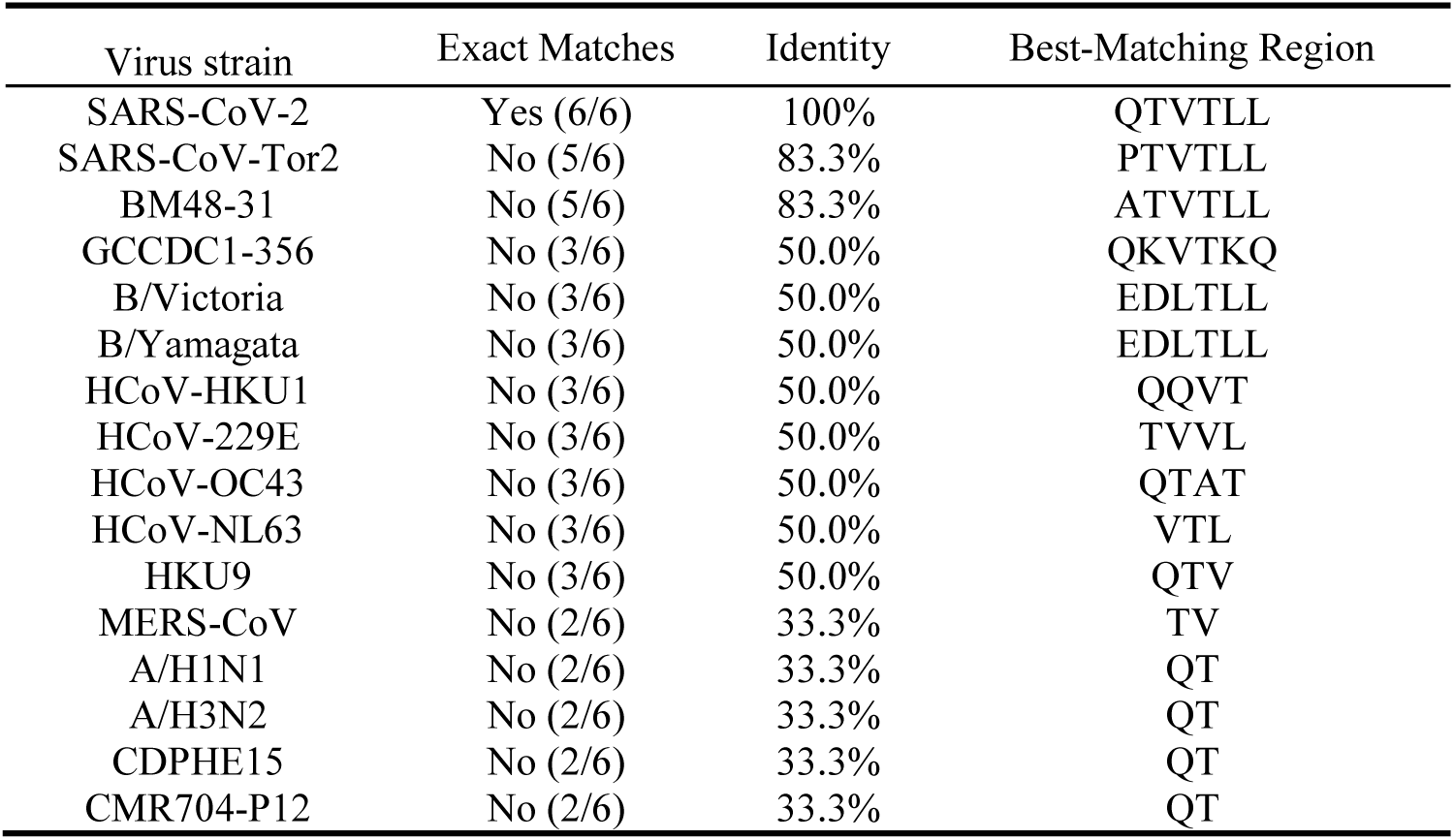
Pairwise local alignment analysis of the ^390^QTVTLL^395^ motif across viral nucleoproteins using EMBOSS WATER. Alignment parameters were as follows: PAM30 substitution matrix, gap opening penalty of 10, and gap extension penalty of 0.5. "Exact matches" indicates the number of residues identical to the ^390^QTVTLL^395^ core motif, which contains six residues. "Identity" denotes the percentage of sequence identity within the best-matching region. "Best-matching region" indicates the amino acid segment produced the highest alignment score. An exact 6/6 match was observed only in SARS-CoV-2; SARS-CoV, the closest relative, retained a single mismatched residue.

### The QTVTLL Sequence Located on the Exposed Surface and Disordered Region of the N Protein

Structural prediction analyses indicated that the ^390^QTVTLL^395^ epitope is located within the flexible, intrinsically disordered C-terminal tail of the N protein, downstream of the C-terminal dimerization domain (Fig 7). Secondary structure analysis showed that this segment was predicted to adopt a predominantly random-coil conformation (Fig 7A and B), and Kyte-Doolittle hydropathy analysis indicated a hydrophilic profile, consistent with potential surface exposure (Fig 7C). ScanProsite domain annotation placed that this region lies within the disordered tail, outside both the N-terminal domain (NTD) and the C-terminal dimerization domain (CTD) (Fig 7D). BepiPred 3.0 analysis assigned this region a linear B-cell epitope propensity score substantially above the default threshold of 0.151 (Fig 7E). The AlphaFold 3 structural model further suggested that the ^390^QTVTLL^395^ region adopts a surface-exposed and accessible conformation in the predicted three-dimensional structure (Fig 7F).

**Fig 7.**
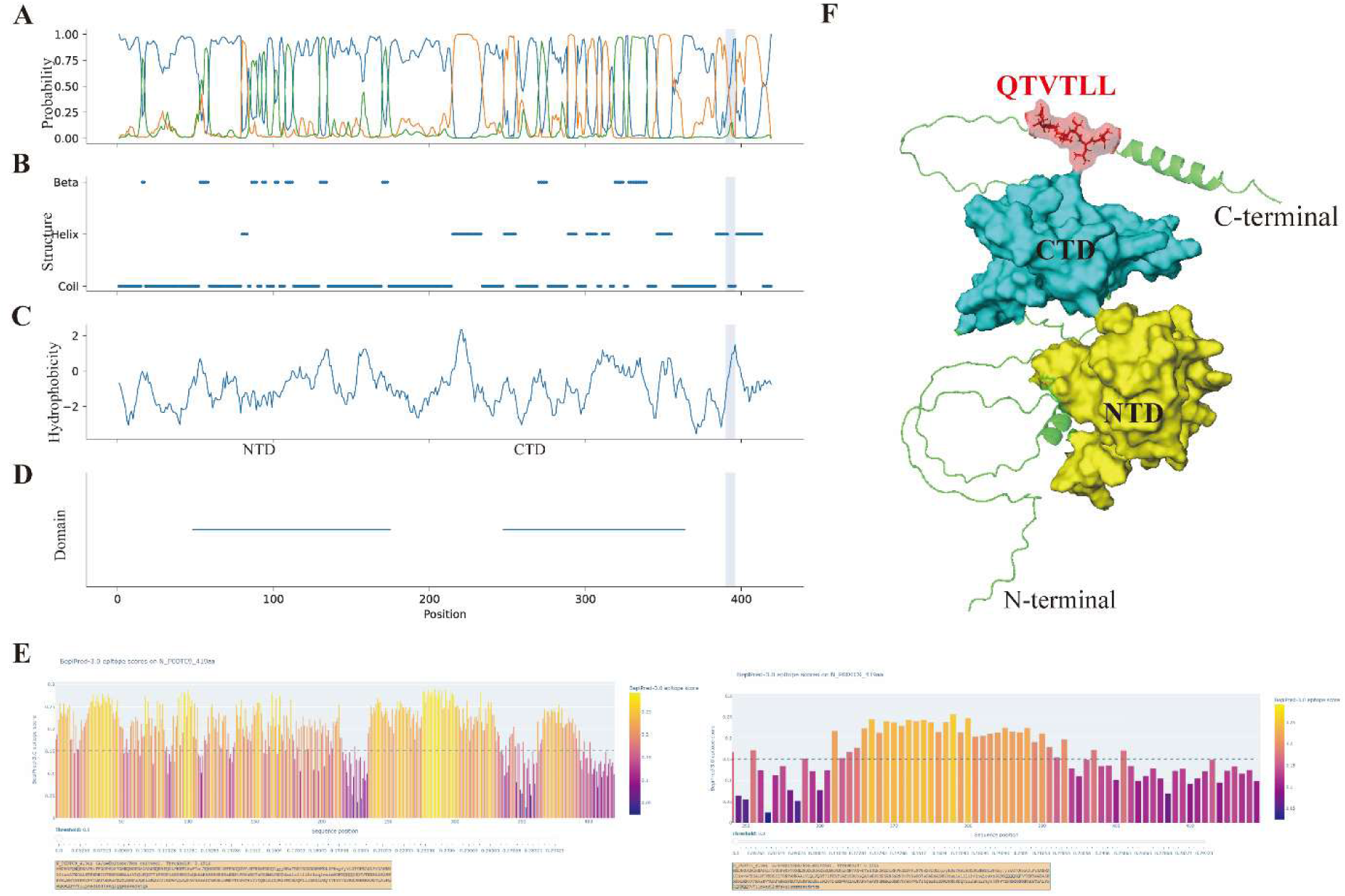
Predicted structural features of the ^390^QTVTLL^395^ epitope region. (A) Secondary structure probability distribution predicted by PSIPRED. (B) Secondary structure state assignment predicted by PSIPRED. (C) Kyte-Doolittle hydropathy plot generated with a window size of 9. (D) ScanProsite domain annotation showing the approximate positions of N-terminal domain (NTD) and C-terminal dimerization domain (CTD) of SARS-CoV-2 N protein. (E) Linear B-cell epitope propensity score predicted by BepiPred 3.0; the dashed line denotes the default threshold of 0.151. (F) AlphaFold3-predicted three-dimensional structural model of the full-length N protein, with the ^390^QTVTLL^395^ region in the flexible C-terminal tail highlighted as a surface-exposed conformation.

### Permission to use copyrighted material

NA

## Discussion

This study identifies ^390^QTVTLL^395^ in the C-terminal tail of the SARS-CoV-2 nucleocapsid (N) protein as the minimal linear epitope required for recognition by mAb N179. This conclusion is supported by sequential truncation analysis combined with Western blotting and ELISA, which together narrowed N179 binding to a compact six-residue motif. These findings define a sequence-dependent antibody-binding determinant rather than a broad C-terminal antigenic region, thereby providing a molecular basis for interpreting the diagnostic performance of the previously developed N179-based colloidal gold immunochromatographic test strip.

The diagnostic relevance of this finding lies in the fact that the performance of antibody-based antigen tests depends on the stability of the actual antibody-binding site, rather than on the overall conservation of the target antigen alone (17). Although the N protein is generally more conserved than the spike protein, specific N-protein substitutions including D399N and R203M, have been reported to impair the sensitivity of antigen detection assays (12, 18). These observations underscore the need to evaluate diagnostic antibodies at the epitope level. To address this issue, we evaluated three features relevant to diagnostic antibody performance: conservation of the N179-binding motif across representative SARS-CoV-2 variants, sequence distinction from related respiratory viruses, and predicted surface accessibility compatible with linear antibody recognition.

The predicted structural features of ^390^QTVTLL^395^ support its accessibility to antibody recognition. Residues 390 to 395 are located downstream of the structured C-terminal dimerization domain, within the flexible C-terminal tail of the N protein (19). Secondary-structure prediction, hydropathy analysis, B-cell epitope propensity evaluation, and AlphaFold-based structural modeling were consistent with a surface-exposed and predominantly flexible region compatible with linear antibody recognition. This structural context is relevant because diagnostic antibodies targeting linear epitopes require access to their binding sites under antigen-detection conditions. The intrinsically disordered nature of the C-terminal tail may allow this short motif to remain accessible across different structural states of the N protein, which may help explain the recognition of recombinant N fragments by mAb N179 in Western blotting and ELISA. However, these structural interpretations should be considered predictive rather than definitive, because the structure of the mAb N179–epitope complex was not experimentally determined.

The identification of ^390^QTVTLL^395^ further refines the antigenic map of the SARS-CoV-2 N-protein C-terminal region. Several linear B-cell epitopes have previously been mapped to this region, including ⁴⁰¹DFSKQLQQ⁴⁰⁸ and ³⁹³TLLPAADLDDFSKQL⁴⁰⁷. These reported epitopes range from 8 to 15 residues and partially overlap with, or lie adjacent to, the motif identified in the present study (20, 21). In comparison, ^390^QTVTLL^395^ represents a minimal six-residue binding unit for mAb N179. The major implication of this finding is not simply that the epitope is shorter, but that mAb N179 recognition can now be assigned to a precise ly delimited motif. This level of resolution allows variant-associated substitutions to be evaluated directly at the actual antibody-binding site rather than across a broader C-terminal antigenic region(15). The observation that deletion of either Q390 or L395 abolished immunoreactivity further indicates that the integrity of the full six-residue motif is required for mAb N179 binding.

The conservation analysis provides important support for the diagnostic relevance of this motif. Multiple sequence alignment of N-protein sequences from 11 representative SARS-CoV-2 lineages, including early variants such as Alpha and Beta as well as Omicron sub-lineages BA.1, BA.2, and BA.3.2, showed that ^390^QTVTLL^395^ was completely conserved across all variants analyzed. This finding is notable because several variants have accumulated substitutions elsewhere in the N protein, and previous reports have shown that certain N-protein mutations can affect antigen-test performance (12, 18). The absence of amino acid changes within the mAb N179-binding motif among the lineages examined suggests that this epitope lies within a locally sequence-stable region of N protein. Whether this stability reflects functional constraint or limited sampling of tolerated substitutions remains to be determined, but the observed conservation provides a plausible explanation for the potential durability of N179-mediated recognition across the variants analyzed.

The sequence-specificity analysis further supports the SARS-CoV-2 specificity of this epitope among the viruses examined. Stringent local pairwise alignment using EMBOSS WATER with a PAM30 substitution matrix showed that a complete 6/6 match to ^390^QTVTLL^395^ was detected only in SARS-CoV-2. No identical continuous six-residue motif was identified in any of the seven other human coronaviruses, four influenza viruses, or five bat coronaviruses examined (22). SARS-CoV and bat coronavirus BM48-31 each contained a single substitution at the first residue of the motif, resulting in a 5/6 match, whereas the remaining viruses showed only partial and discontinuous similarity in their best matching regions. These data indicate that ^390^QTVTLL^395^ is conserved among the SARS-CoV-2 variants analyzed and distinct from the corresponding regions of the other respiratory viruses examined. Although sequence analysis alone cannot exclude cross-reactivity against every possible respiratory virus or naturally occurring isolate, these findings provide a plausible sequence-level explanation for the previously observed lack of detectable cross-reactivity of the N179-based immunochromatographic assay with common respiratory pathogens.

This study has several limitations. First, the sequential truncation strategy enabled precise localization of the minimal linear epitope but would not identify conformational determinants that depend on the three-dimensional structure of the antigen (23). Second, although computational analyses supported the predicted accessibility of ^390^QTVTLL^395^, the structure of the N179-epitope complex was not experimentally resolved, and the atomic contacts responsible for antibody recognition remain unknown. Third, the kinetic parameters of the N179–epitope interaction were not measured by surface plasmon resonance, biolayer interferometry, or related biophysical methods (24). Fourth, although deletion analysis defined the minimal motif, site-directed mutagenesis or alanine scanning of individual residues within ^390^QTVTLL^395^ would further clarify the contribution of each residue to N179 binding (25). Finally, the sequence analysis included representative SARS-CoV-2 lineages and selected related respiratory viruses, but broader surveillance-scale analysis would be useful for monitoring rare or emerging substitutions within this motif. Future studies combining residue-level mutagenesis, kinetic binding assays, structural analysis, and testing against recombinant N proteins or clinical samples carrying relevant N mutations would further define the diagnostic robustness of this epitope.

In summary, this work maps ^390^QTVTLL^395^ as a conserved, predicted surface-accessible, and SARS-CoV-2-specific linear epitope recognized by mAb N179. By integrating fine epitope mapping, cross-variant conservation analysis, sequence-specificity evaluation, and structural prediction, this study provides an epitope-level explanation for the diagnostic performance and specificity profile of the N179-based assay. These findings support ^390^QTVTLL^395^ as a rational reference motif for diagnostic antibody selection, assay revalidation, and epitope-informed optimization of SARS-CoV-2 antigen detection strategies.

## Acknowledgments

The authors thank all members in biotechnology laboratory of Zhuhai Campus of Zunyi Medical University for technical assistance and helpful discussions.

## Data availability statement

The raw data supporting the findings of this study are available from the corresponding author upon reasonable request. The SARS-CoV-2 N protein sequences used for alignment are publicly accessible in the NCBI database under the accession numbers detailed in section 2.4.1. The AlphaFold structural model is available via AlphaFold DB accession number AF-0000000365840321-v1.

## Clinical trials

NA

## Ethics approval

NA

